# Chronic inflammation delays cell migration to villi in the intestinal epithelium

**DOI:** 10.1101/248344

**Authors:** Daniele Muraro, Aimee Parker, Laura Vaux, Sarah Filippi, Alexander G. Fletcher, Alastair J. Watson, Carmen Pin, Philip K. Maini, Helen M. Byrne

## Abstract

The intestinal epithelium is a single layer of cells which provides the first line of defence of the intestinal mucosa to bacterial infection. Cohesion of this physical barrier is supported by renewal of epithelial stem cells, residing in invaginations called crypts, and by crypt cell migration onto protrusions called villi; dysregulation of such mechanism may render the gut susceptible to chronic inflammation. The impact that excessive or misplaced epithelial cell death may have on villus cell migration is currently unknown. We integrated cell-tracking methods with computational models to determine how epithelial homeostasis is affected by acute and chronic inflammatory cell death. Parameter inference reveals that acute inflammatory cell death has a transient effect on epithelial cell dynamics, whereas cell death caused by chronic inflammation causes a delay in the accumulation of labelled cells onto the villus compared to control. Such a delay may be reproduced by using a cell-based model to simulate the dynamics of each cell in a crypt-villus geometry, showing that a prolonged increase in cell death slows the migration of cells from the crypt to the villus. This investigation highlights which injuries (acute or chronic) may be regenerated and which cause disruption of healthy epithelial homeostasis.

## Introduction

The intestinal epithelium is a rapidly self-renewing tissue, formed of a single layer of cells, that covers the luminal surface of the small and large intestine, providing a barrier to bacterial infection. Epithelial cells in the small intestine are organised into numerous protrusions, termed villi, and invaginations, termed crypts of Lieberkühn. Self-renewal is sustained by the proliferative activity of adult stem cells at the base of intestinal crypts whose progeny proliferate and then differentiate into the functionally distinct epithelial subtypes that migrate onto the villus where they are eventually shed into the gut lumen [1]. Such cellular dynamics can be thought of as a ‘conveyor belt’ where cell proliferation acts as the principal driving force for cell migration on villi [2]. Perturbations of this tightly controlled process may be responsible for the development of serious diseases. For example, excessive or misplaced cell death may disrupt barrier function and cause chronic inflammation; on the other hand, deficiency in cell death may lead to cancer development [3]. In combination with experimental studies, mathematical modelling helps us to disentangle the complex interactions underlying the self-renewal of the intestinal epithelium under healthy and pathological conditions. The cellular dynamics of the intestinal epithelium have been studied using a variety of theoretical approaches, including compartmental models based on ordinary differential equations (ODEs) [4], [5], continuum models [6], cell-based models [7], [8], [9] - [17]. Experimental and theoretical studies of the influence of reduced or halted proliferation on epithelial homeostasis showed a pronounced coupling of cell proliferation with cell migration onto villi [18], [2]. However, it remains unknown whether an increase in cell death in the epithelium affects villus cell migration and how excessive cell death on a particular villus influences epithelial homeostasis in neighbouring crypts.

Here, we use a multidisciplinary approach to determine how two types of induced inflammatory cell death affect cell migration on villi in two regions of the mouse small intestine (ileum and duodenum). Inflammatory cell death was induced by exposure to TNF*α*, a cytokine involved in systemic inflammation, for different time periods and levels. During ‘acute inflammation’ the mice expressed a high level of TNF*α* for about 90 minutes before it dropped back to the baseline level; this treatment caused cell death and detachment from villus tips. During ‘chronic inflammation’ the mice expressed a lower dose of TNF*α* for two weeks continuously prior to and during the measurements; this treatment induced less severe, but more persistent, cell death.

To investigate epithelial cell dynamics during acute and chronic inflammation, we applied cell-tracking methods to monitor accumulation of labelled cells along the crypt-villus axis following exposure of healthy crypts to high and low doses of TNF*α*. We generated experimental time courses from crypt-villus epithelial units (CVEU) indicating the number of cells that were tracked (labelled) in the crypt and villus com-partments. We then derived complementary information from two different computational approaches, namely cell-based and compartmental models, as follows. We replicated the experiments, simulating injury in a cell-based model in which cells are confined to a 2D surface comprising four crypts, adjacent to a villus, with cells moving according to a nearest-neighbour-defined repulsive force [15]. This model allows us to describe the spatial dynamics of cells on a crypt-villus geometry and to generate simulated time-courses that can be compared to the experimental time courses via time-dependent compartmental models as described below; however, the parameters of the cell-based model cannot be easily inferred from the experimental data since fitting such a detailed and stochastic model would be computationally intensive. By contrast, the compartmental models are described by a smaller number of parameters, since they do not account for the parameters associated with the crypt-villus geometry, and their simulation is computationally inexpensive. These advantages come at the expense of the biological detail included in the model: it accounts only for the time evolution of the number of cells in the crypt and villus compartments and neglects spatial effects. A schematic of our approach is presented in Figure 1. As in Parker et al [2], we first developed a compartmental model that distinguishes two compartments, crypt and villus, and obtained quantitative estimates of parameters describing cell proliferation, migration and death by fitting it to the experimental data using a variant of Hamiltonian Monte Carlo (the No-U-Turn sampler) [19]. The posterior predictive distributions, showing the simulated time evolution of the number of labelled cells in the crypts and in the villi, produced fits that are in good agreement with the trend of the experimental time courses and highlighted that chronic inflammation caused an increase in cell death, which, in turn, generated a decrease in the accumulation of labelled cells on villi. By contrast, acute inflammation generated a similar, but small, delay. The two-compartment model relies on the simplifying assumption that all cells in the crypts proliferate, whereas in practice only some of them do. For this purpose, we extended the two-compartment model by including a further compartment which enables us to distinguish between proliferative and non-proliferative crypt cells. As for the two-compartment model, the three-compartment model produced fits that are in good agreement with the experimental time courses; in addition, it generated predictions about the dynamics of the number of proliferative and non-proliferative cells in the crypt. To investigate how an increase in cell death may influence an accumulation of labelled cells from the crypt to the villus, we then used the cell-based model to simulate injury due to treatments causing acute and chronic inflammation. Quantitative estimates of the parameters of the compartmental models, derived by model fitting against these synthetic time courses, revealed a decrease in the accumulation of labelled cells on villi under chronic injury and a minor decrease under acute injury, as experimentally observed. The consensus between the compartmental and cell-based models suggests that injuries caused by acute and chronic inflammation manifest themselves via treatment-specific decreases in the accumulation of labelled cells on villi.

**Figure 1:**
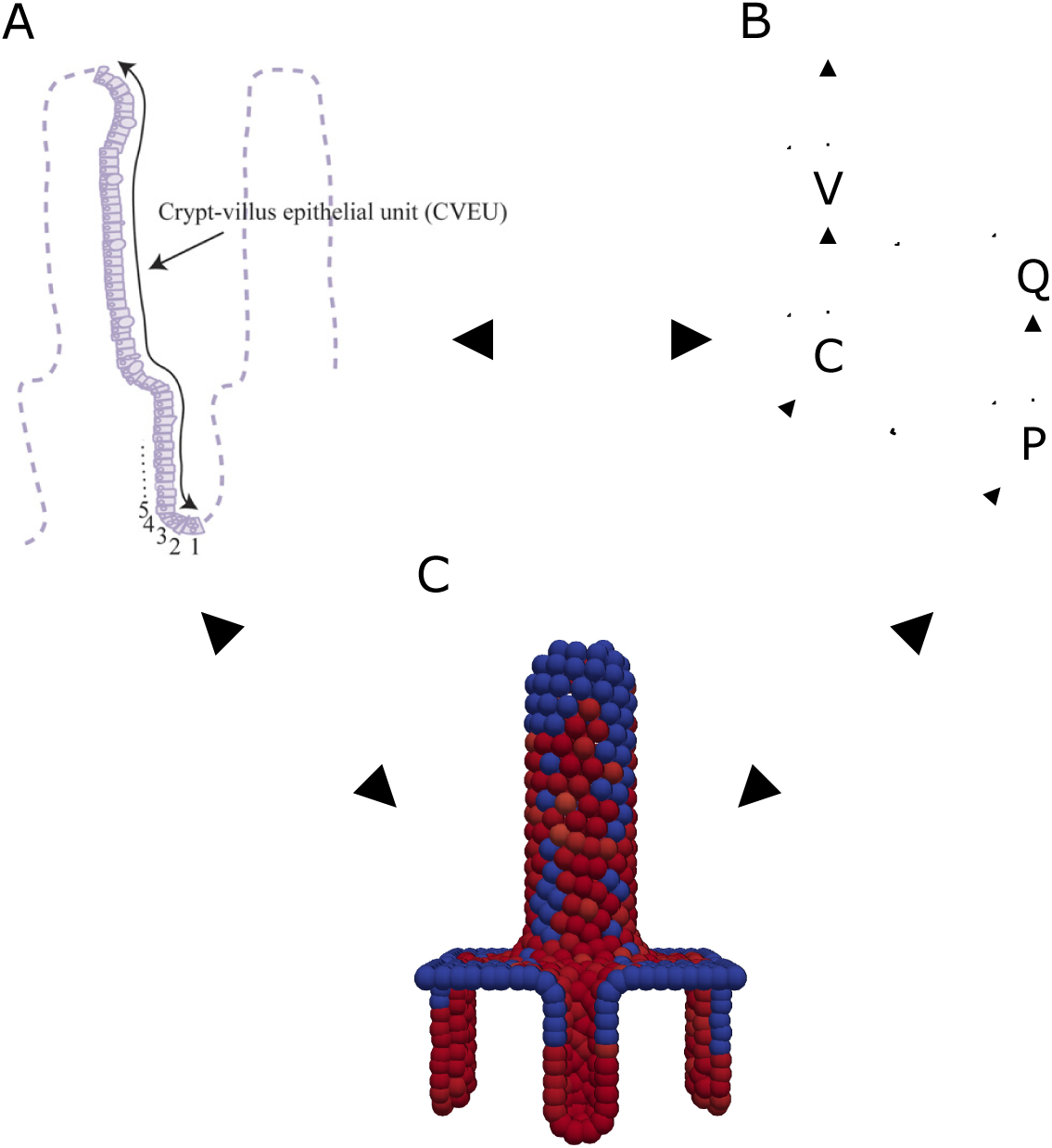
Schematic of our approach. A) Experimental time courses are derived from crypt-villus epithelial units (CVEU) and are analysed by counting labelled cells in the crypt and villus compartments. B) A compartment-based model accounting for crypt cells (C) and villus cells (V) allows us to quantify cell migration under injury and control. A model extension which distinguishes between a proliferative (P) and a non-proliferative (Q) compartment generates predictions on the dynamics of the number of proliferative and non-proliferative cells in the crypt. C) A multi-cellular model allows for replication of the experiment and for generation of simulated data; in red and blue are presented labelled and unlabelled cells, respectively. The arrows are interpreted as follows: A → B (experimental data informing model parameterisation): The experimental data allow for inference of the compartment-based model parameters; B → A (model prediction): The posterior predictive distributions highlight a decrease in the accumulation of labelled cells on villi which is specific of the type of injury (acute or chronic); A → C (experimental background informing model development): The injuries caused by acute inflammation (death and detachment of cells from villus tips) and chronic inflammation (less severe, but more persistent, rates of cell death) inform the replication of the experiments (simulated injuries) by means of cell-based simulations and allow for generation of simulated time courses; B ↔ C (consensus between models): The posterior predictive distributions obtained when fitting the compartment based models to data simulated by means of the cell-based model highlight a qualitative agreement with the fits to the experimental data; C → A (model prediction): The consensus between the models supports the driving role of the injuries caused by acute and chronic inflammation in generating treatment specific decrease in the accumulation of labelled cells on villi.

## Methods

### Experimental data

#### Animals

All animal experiments were conducted in strict accordance with the Home Office Animals (Scientific Procedures) Act 1986. Female C57BL/6 mice, aged 10-12 weeks and weighing at least 20 g prior to use in experiments, were housed and maintained in SPF conditions at the University of East Anglia, Norwich, UK in accordance with HO regulations, and all procedures were performed by fully-trained and licenced researchers. Experimental animals were closely monitored and were sacrificed by rising CO2 and cervical dislocation, at the timepoints described in the text, prior to subsequent tissue collection. All animals were regularly monitored for clinical signs, any displaying signs beyond those expected within the moderate limits of the procedures would be immediately sacrificed by the above methods and not included in experimental data.

#### Induction of inflammation, cell labelling and tissue processing

Transient, acute inflammation was induced by single intraperitoneal injection of recombinant murine TNF*α* (Peprotech, London, UK) at 0.5 mg/kg. Chronic inflammation was achieved by hydrodynamic tail vein delivery of TNF*α*-expressing plasmid (originally a kind gift from C. Gunther, Erlangen, Germany). TNF*α* overexpression was confirmed by specific ELISA (Thermo Fisher Scientific, Waltham, USA) for elevated levels in blood plasma over a minimum of 14 days, and in liver and intestinal tissue lysates post mortem. The thymine analogue 5-bromo-2-deoxyuridine, BrdU, (Sigma-Aldrich, Paisley, UK) was administered at 50 mg/kg body weight by single intraperitoneal injection. In the case of acute inflammation, BrdU was delivered simultaneously with TNF*α*. In the chronic inflammation experiments, BrdU time-courses were performed once elevated blood TNF*α* levels had been established. At time points from 1h - 48h post BrdU-administration, mice were euthanised and intestinal tracts were removed, dissected, formalin-fixed and paraffin embedded. Transverse sections of duodenum and ileum were prepared at 5*μm* and were immunostained for BrdU using biotinylated anti-BrdU antibody (AbCam, Cambridge, UK), Neutravidin-HRP (Thermo Fisher), and diaminobenzidine reaction (DAB, Dako, Glostrup, Denmark). Villus cell shedding was confirmed histologically by Caspase-3 (anti-CC3, R&D Systems, Minneapolis, USA) labelling of apoptotic cells in FFPE duodenal and ileal sections counterstained with H&E.

#### Data collection

Collection of the experimental dataset followed the format described in Parker et al [2]. The number of unlabelled and BrdU-labelled cells by position, from crypt base to neighbouring villus tip, was counted for 30-50 individual hemi crypt-villus units per tissue section per mouse, with counts recorded as binary values. This generated, for each replicate and at each time point, a binary vector whose length varied with the particular sample. Counts were taken at multiple time-points post-delivery of BrdU and post induction of inflammation. The counts and the code to calculate the experimental time courses are reported in the Supplementary Data (folder Counts at https://tinyurl.com/y9xk3nsk). The number of samples for each time point are shown in Supplementary Tables 1 and 2. The boundary between the crypt and villus compartment was estimated from all datasets obtained during the first 2 hours after BrdU injection as the cell position closest to the crypt bottom and such that the fraction of labelled cells in the villus is smaller than 0.01. The time courses obtained from the ileum are presented in Figure 2a; the corresponding time courses from the duodenum are presented in Figure 2b.

**Figure 2:**
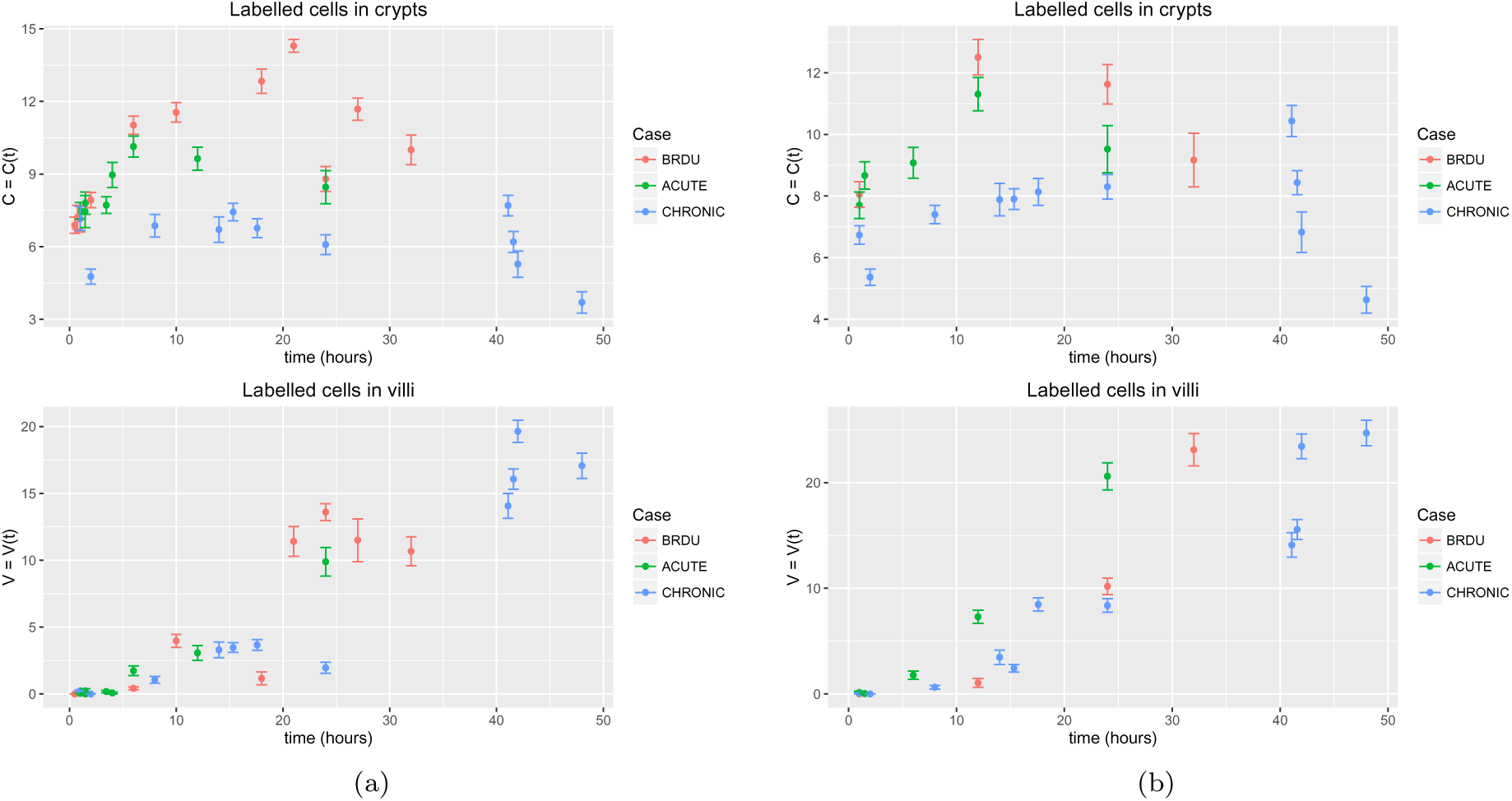
Experimental data. Time series representing average numbers of cells in crypts *C* = *C*(*t*) and villi *V* = *V*(*t*) during acute inflammation, chronic inflammation and control (BrdU) in ileum (a) and duodenum (b). Error bars indicate standard errors.

#### Compartment-based models

To analyse the spatio-temporal dynamics of BrdU-labelled cells we derived two compartmental models formulated as a system of time-dependent ordinary differential equations (ODEs). The first model treats the crypt-villus unit as two distinct compartments and distinguishes the cell numbers in the crypt and villus; the second model decomposes the crypt-villus unit into three compartments and distinguishes between proliferative and non-proliferative cells in the crypt. For simplicity, and to allow for parameter estimation, in what follows we model labelled cells only.

#### Two-compartment model

We distinguish two cellular compartments: labelled cells in the crypt, whose number at time t is denoted by *C* = *C*(*t*), and labelled cells in the villus, whose number is denoted by *V* = *V*(*t*). We introduce two parameter thresholds *C*\*, *V*\* such that when *C*(*t*) > *C*\* labelled cells in the crypt start migrating onto the villus, and when *V*(*t*) > *V*\* cells begin to be shed from the villus. We denote condition-specific death rates in the crypt and villus compartment as follows:

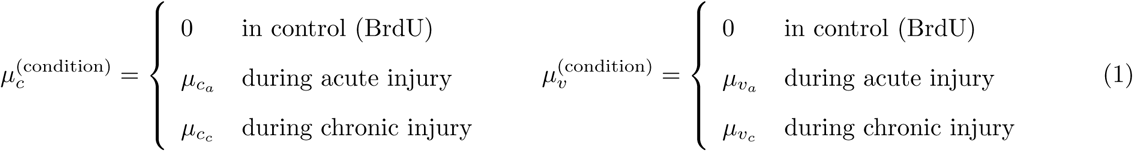

where *μ*_*c*_*a*__, *μ*_*c*_*c*__, *μ*_*v*_*a*__, *μ*_*v*_*c*__ are positive constants. The two-compartment model is described by the following pair of ODEs:

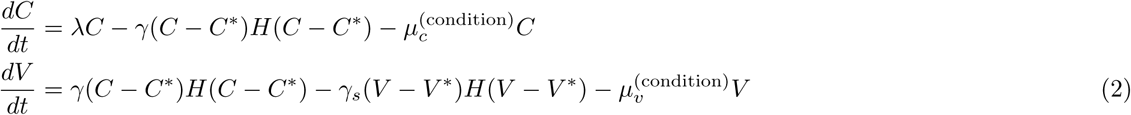

where *λ* is the net cell proliferation rate (cell proliferation minus cell death rate), *γ* is the cell migration rate between the two compartments, *γ*_*s*_ is the cell shedding rate from the villus, *H* is the Heaviside function:

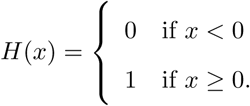

Model parameters and initial conditions, included in the set of parameters to be estimated, are listed in Table 1.

**Table 1:** Summary of the parameters and initial conditions that appear in the two-compartment model defined by Equations (2).

| Parameter | Description | Units |
| --- | --- | --- |
| $\lambda$ | Proliferation rate | $h^{-1}$ |
| $\gamma$ | Migration rate into the villus | $h^{-1}$ |
| $\mu_{c_a}$ | Death rate in the crypt during acute injury | $h^{-1}$ |
| $\mu_{v_a}$ | Death rate in the villus during acute injury | $h^{-1}$ |
| $\mu_{c_c}$ | Death rate in the crypt during chronic injury | $h^{-1}$ |
| $\mu_{v_c}$ | Death rate in the villus during chronic injury | $h^{-1}$ |
| $C^*$ | Number of labelled cells in the crypt<br>above which migration to the villus starts | - |
| $C_0 \equiv C(t=0)$ | Initial number of labelled cells in crypt | - |
| $V_0 \equiv V(t=0)$ | Initial number of labelled cells on villus | - |

#### Three-compartment model

The three-compartment model subdivides the crypt into proliferative and non-proliferative cells and defines the following compartments: labelled proliferative cells in the crypt, whose number at time *t* is denoted by *P* = *P*(*t*); labelled non-proliferative cells in the crypt, whose number is denoted by *Q* = *Q*(*t*) and labelled non-proliferative cells on the villus, whose number is denoted by *V* = *V*(*t*) (cells on the villus do not proliferate). For comparison with the two compartment model, we also denote by *C*(*t*) = *P*(*t*) + *Q*(*t*) the total number of labelled cells in the crypt. We introduce three parameter thresholds *P*\*, *Q*\*, *V*\* such that when *P*(*t*) > *P*\* labelled proliferative cells start migrating onto the villus, when *Q*(*t*) > *Q*\* non-proliferative labelled cells start migrating onto the villus and when *V*(*t*) > *V*\* cell shedding begins to occur from the villus. Both P and Q cells can migrate onto villi and are affected by acute and chronic inflammation; for simplicity, we assume equal rates of cell transfer onto villi and of cell death in the crypts. We denote condition specific death rates in the crypt and villus compartments as for the two-compartment model (see Equations (1)). The three-compartment model is described by the following system of time-dependent ODEs:

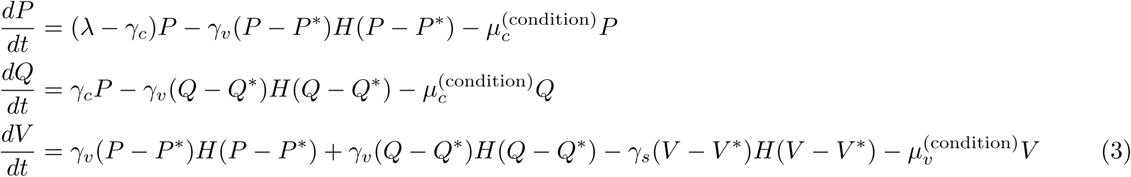

where *λ* is the cell net proliferation rate, *γ*_*c*_ is the rate at which cells differentiate from a proliferative to a non-proliferative state, *γ*_*v*_ is the rate at which cells migrate onto the villus, *γ*_*s*_ is the rate of cell shedding from the villus and *H* is the Heaviside function. Model parameters and initial conditions, included in the set of parameters to be estimated, are listed in Table 2.

**Table 2:** Summary of the parameters and initial conditions that appear in the three-compartment model defined by Equations (3).

| Parameter | Description | Units |
| --- | --- | --- |
| $\lambda$ | Proliferation rate | $h^{-1}$ |
| $\gamma_c$ | Migration rate from proliferative to non-proliferative state | $h^{-1}$ |
| $\gamma_v$ | Migration rate into the villus | $h^{-1}$ |
| $\gamma_s$ | Cell shedding rate | $h^{-1}$ |
| $\mu_{c_a}$ | Death rate in the crypt during acute injury | $h^{-1}$ |
| $\mu_{v_a}$ | Death rate in the villus during acute injury | $h^{-1}$ |
| $\mu_{c_c}$ | Death rate in the crypt during chronic injury | $h^{-1}$ |
| $\mu_{v_c}$ | Death rate in the villus during chronic injury | $h^{-1}$ |
| $P^*$ | Number of proliferative and labelled cells in the crypt<br>above which migration to non-proliferative state starts | - |
| $Q^*$ | Number of non-proliferative and labelled cells in the crypt<br>above which migration to the villus starts | - |
| $V^*$ | Number of labelled cells in the villus<br>above which cell shedding starts | - |
| $P_0 \equiv P(t=0)$ | Initial number of labelled proliferative cells in crypt | - |
| $Q_0 \equiv Q(t=0)$ | Initial number of labelled non-proliferative cells in crypt | - |
| $V_0 \equiv V(t=0)$ | Number of labelled cells in villus (all non-proliferative) | - |

#### Cell-based simulations

We simulated injury by using a cell-based simulation of cell dynamics on a patch of intestinal epithelium composed of multiple crypts and a single villus, previously developed by Mirams et al. [15], and generated synthetic time courses.

The model is a stochastic 3D off-lattice cell centre based model confined to a 2D surface comprising four crypts that surround a single villus; the crypts and the villus are modelled using a cylindrical geometry with spherical rims. Cell movement is driven by a nearest-neighbour-defined force, previously employed by Meineke et al. [9]. Each pair of neighbouring nodes is assumed to be connected by a linear spring. The force of node *i* is given by

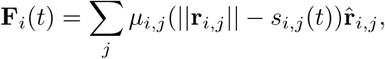

where *μ*_*i*, *j*_ is the spring constant for the spring between nodes *i* and *j*, *s*_*i*, *j*_ (*t*) is its natural length at time *t*, r_*i*,*j*_ is their relative displacement and a hat (^∧^) denotes a unit vector. Cells moving above a plane defined at the villus tip are removed due to anoikis. Injury is simulated either by removing cells randomly (during chronic injury) or by initially removing cells that are above a plane defined at ⅔ of the villus height (to account for the experimentally observed detachment of cells at the top third of the villus during acute injury). Cell proliferation depends on a decreasing gradient of the Wnt family of morphogens from the crypt to the villus [21] and it is modelled by defining a linear gradient in Wnt concentration up the crypt, allowing cells to divide when their Wnt concentration exceeds a fixed threshold (see SimpleWntCellCycleModel class for details [15]).

All simulations were initialised without including any random removal of cells and were run for 1000 h with default parameter values [15], at which time the total number of cells in the crypts and in the villus was approximately constant. After initialisation, cell-based simulations at homeostasis (control) were run for 80 hours with default parameter values. During this time period, crypt cells were labelled and their lineage was tracked according to their ancestor proliferative cell. Acute injury was modelled by initially detaching cells from the top third of the villus. Regeneration of this area, due to cell migration from the crypts, was simulated for 80 hours. Chronic injury was introduced by randomly killing cells in the crypts and the villus with the default probability value *p* = 0.005 *h*^‒1^.

#### Parameter estimation

The compartmental models were solved using the R-packages deSolve (Classical Runge-Kutta 4th Order Integration) [20] and RSTAN [19]. STAN is a C++ library that performs Bayesian inferences using the No-U-Turn sampler (a variant of Hamiltonian Monte Carlo); the RSTAN package conveniently allows STAN to be used from R. RSTAN was applied to Equations (2) and (3) with uniform priors represented in Supplementary Figures 13, 15, 17, 19, 21, 23. Convergence diagnostics were then calculated for four Markov Chain Monte Carlo (MCMC) chains using the R package CODA, which provides routines for output analysis and diagnostics for MCMC [22]. Where multi-modality was highlighted by chains mixing around different modes, the chains with the highest fit quality (STAN log probability variable lp_—_) were selected. The initial conditions for *P*(*t*), *Q*(*t*) (three-compartment model) and *C*(*t*) (two-compartment model) were included in the set of parameters to be estimated by applying MCMC. Since the number of labelled cells in the villus (*V*(*t*)) is approximately zero at the start of the time courses, we assumed that migration of labelled cells onto villi may be initially neglected.

## Results

In what follows, we first describe the predictions of our compartmental and cell-based models regarding the influence of inflammation on epithelial homeostasis; we then discuss the parameters inferred when fitting the compartmental models to experimental and simulated data.

### Accumulation of labelled cells on villi is delayed during chronic inflammation

The two‐ and three-compartment models were fitted against the experimental data derived from the ileum and the duodenum as described in the Methods section. The resulting posterior predictive distributions are shown in Figures 3, 4 and Supplementary Figures 2, 3. Both models reproduce the trend of the experimental data and show a delay in migration during chronic inflammation compared to control (see Figure 5 and Supplementary Figure 4). Acute inflammation causes a modest delay in cell migration terms in the ileum and a very small decrease in the duodenum (Figure 5 and Supplementary Figure 4).

**Figure 3:**
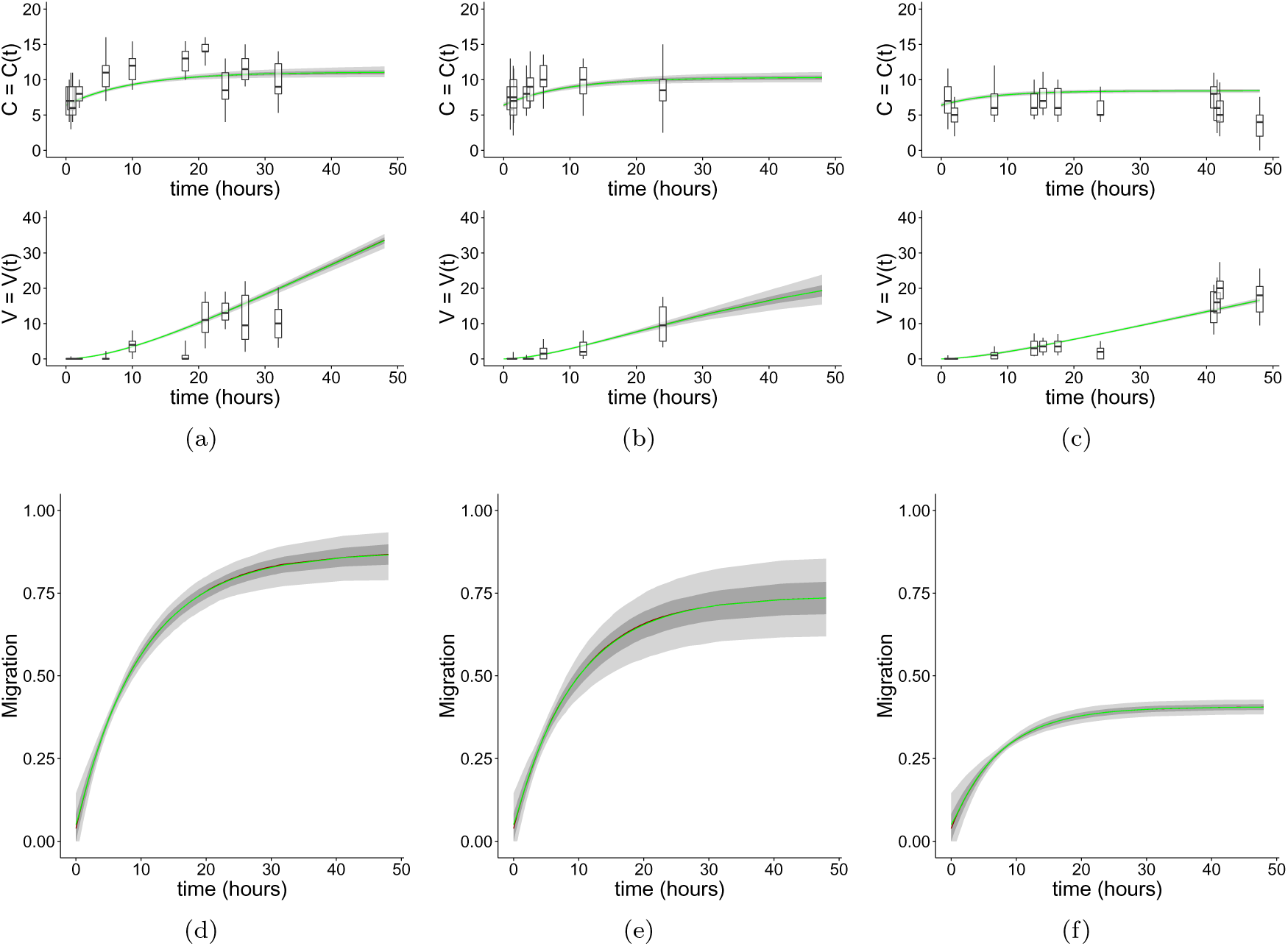
Fits of the two-compartment model to ileal time course data. (a)-(b)-(c) Posterior predictive distributions and estimates of parameter uncertainty obtained by fitting the two-compartment model (Eqs. (1)) against ileal time courses. Posterior predictive distributions inferred from (a) BrdU, (b) acute inflammation, (c) chronic inflammation experimental time courses. Boxplots represent the 0.05, 0.25, 0.75, 0.95 quantiles of the experimental data. Dark and light grey area plots represent the [0.25,0.75] and the [0.25,0.75] quantiles of the posterior predictive distributions, respectively. The green line indicates the posterior mean; the red line, partially overlapping the green line, represents the posterior median. (d)-(e)-(f) Plots representing the posterior predictive distribution of the migration term *γ*(*C* – *C*\*)*H*(*C* – *C*\*) in the ileum obtained from control (BrdU) (d), acute inflammation (e) and chronic inflammation (f) time courses. The contribution to migration is reduced during chronic inflammation.

**Figure 4:**
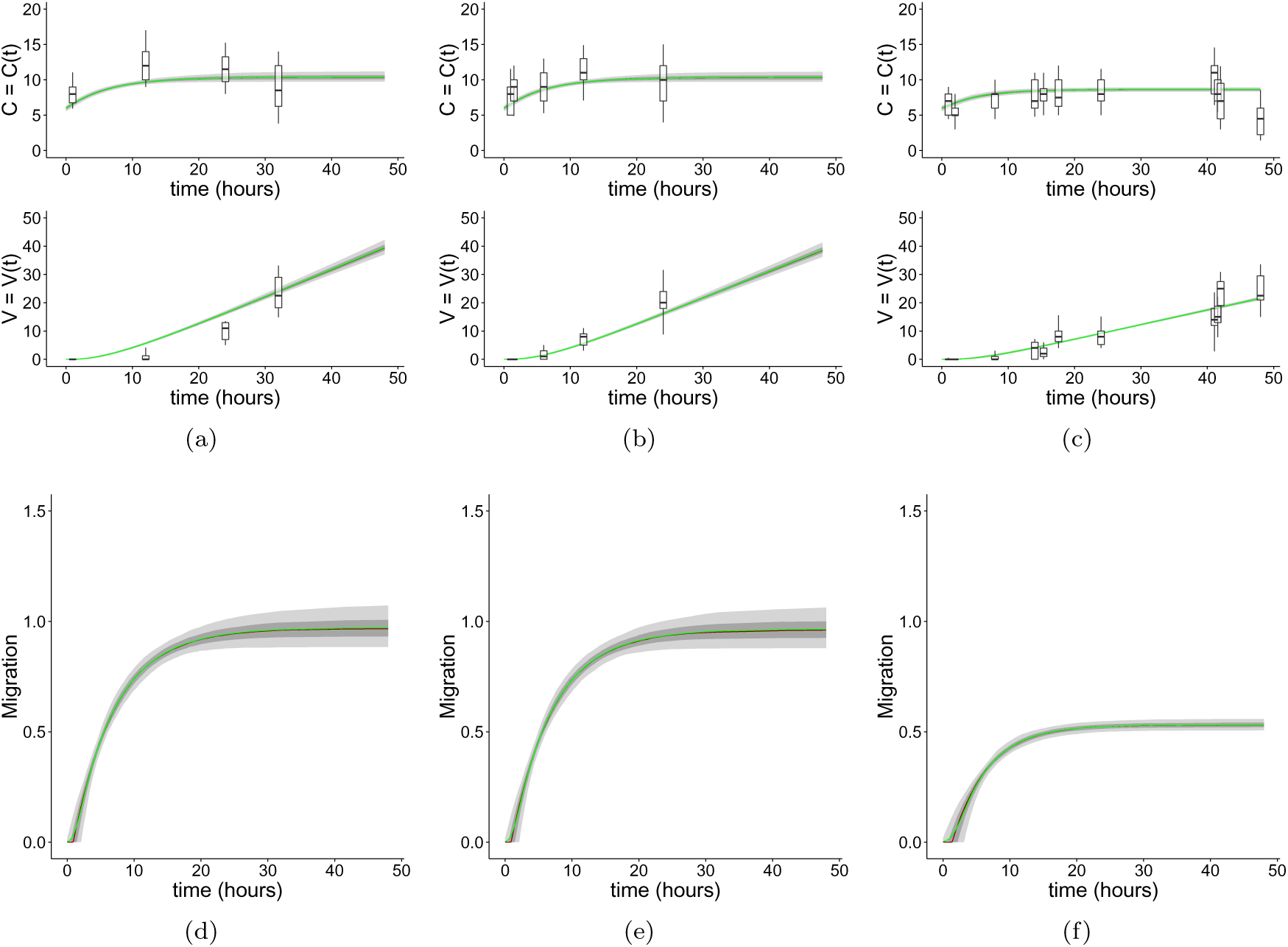
Fits of the two-compartment model to duodenal time course data. (a)-(b)-(c) Posterior predictive distributions and estimates of parameter uncertainty obtained by fitting the two-compartment model (Eqs. (1)) against duodenal time courses. Posterior predictive distributions inferred from (a) BrdU, (b) acute inflammation, (c) chronic inflammation experimental time courses. Boxplots represent the 0.05, 0.25, 0.75, 0.95 quantiles of the experimental data. Dark and light grey area plots represent the [0.25, 0.75] and the [0.25,0.75] quantiles of the posterior predictive distributions, respectively. The green line indicates the posterior mean; the red line, partially overlapping the green line, represents the posterior median. (d)-(e)-(f) Plots representing the posterior predictive distribution of the migration term *γ*(*C* – *C*\*)*H*(*C* – *C*\*) in the duodenum obtained from control (BrdU) (d), acute inflammation (e) and chronic inflammation (f) time courses. The contribution to migration is reduced during chronic inflammation.

**Figure 5:**
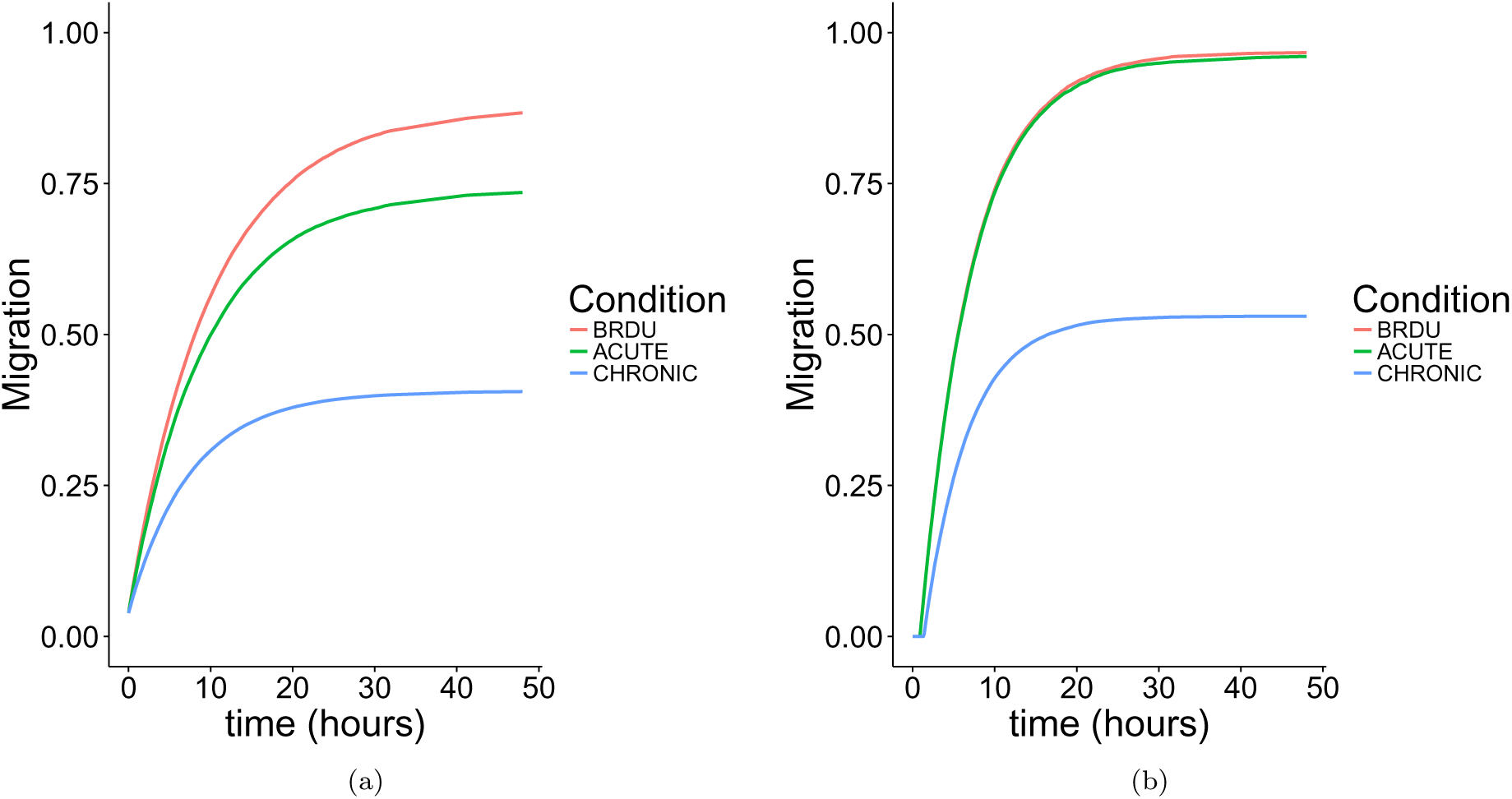
Migration terms of the two-compartment model when fitted against experimental time courses. Plots representing the medians of the posterior predictive distribution of the migration terms *γ*(*C* – *C*\*)*H*(*C* – *C*\*) in ileum (a) and in duodenum (b). Contribution to migration is reduced during simulated chronic inflammation.

### Cell-based simulations suggest that injuries drive treatment specific delays in cell migration

The cell-based model was simulated as described in the Methods section ‘Cell-based simulations’. Typical simulation results are presented in the Supplementary Data (file Cell_Based_Simulations.pptx at https://tinyurl.com/y9xk3nsk). Supplementary Figure 1 shows the mean and standard errors of simulated time series of labelled cells generated by ten simulations for each condition. Compared to control simulations, the simulated persistent, increased rate of cell death associated with chronic injury appears to hinder the migration of labelled crypt cells onto the villus. Conversely, the initial detachment of cells from the villus tip caused by simulated acute injury does not seem to affect significantly cell migration from the crypts to the villus and the villus tip regenerates due to cell migration from the crypts. To confirm these effects, we then fitted the compartmental models to the time courses generated from simulations of the cell-based model. The simulated data were fitted up to 50 hours to emulate the duration of the experimental time courses. Supplementary Figures 5 and 6 show the posterior predictive distributions of the two‐ and three-compartment models together with the predicted migration terms. As observed when applying the model to the experimental data, an increase in cell death caused a delay in the accumulation of labelled cells on villi during simulated chronic inflammation and a minor delay due to simulated acute inflammation (Supplementary Figure 7). The posterior distributions of the parameter are presented in Supplementary Figures 8-23. A prolonged delay in the accumulation of labelled cells on villi during chronic inflammation compared to acute inflammation is caused by the combined increase in the death rates in the crypts (*μ*_*c*_) and in the villi (*μ*_*v*_), (Supplementary Figures 8-11).

### Small regions of the parameter search space allow for good quality fits

Highly correlated parameters may be found in both of the compartment models. In particular, the pairs (*λ*, *μ*_*c*_*c*__) and (*γ*, *C*\*) are the most strongly correlated parameters in the two-compartment model in both tissues (Supplementary Figures 12 and 14); whereas, (*λ*, *γ*_*c*_), (*λ*, *P*\*), (*γ*, *P*\*), (*λ*, *P*_0_), (*P*, *P*^0^), (*P*_0_, *Q*_0_) are the most correlated pairs in the three-compartment model in both tissues (Supplementary Figures 16 and 18). Highly correlated pairs were also found when fitting the compartment models against simulated data; for example, (*λ*, *C*_0_), (*γ*, *C*\*), (*λ*, *γ*_*s*_) in the two-compartment model (Supplementary Figure 20) and (*λ*, *γ*_*c*_), (*λ*, *P*\*), (*γ*_*c*_, *P*\*), (*λ*, *P*_0_), (*P*\*, *P*^0^), (*P*_0_, *Q*_0_) and others in the three-compartment model (Supplementary Figure 22). Notwithstanding this dependence between different parameters, density plots of the posterior distributions highlight that relatively small regions of the parameter search space, defined by uniform prior distributions, allow for good quality fits (Supplementary Figures 13, 15, 17, 19, 21, 23).

### The time thresholds associated with cell migration and cell shedding are most sensitive to crypt parameter

Because of the increase of the death rates *μ*_*c*_ and *μ*_*v*_ during acute and chronic inflammation, we analysed how changes in these parameters in the two-compartment model may affect the time thresholds above which cell migration and cell shedding begin. More precisely, we denoted by 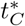 and 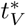 the time thresholds after which *C*(*t*) > *C*\* and *V*(*t*) > *V*\*, respectively. Simulation of the perturbed model highlighted that increasing *μ*_*c*_ causes a delay in both time thresholds, whereas *μ*_*v*_ only affects 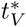 (Supplementary Figure 24). Supplementary Figures 25 - 27 show how the time thresholds vary when all model parameters are varied and highlight that both thresholds are most sensitive to *λ*, *μ*_*c*_ and *C*_0_. Similar effects were found when simulating the three-compartment model by defining the thresholds 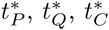 and 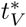, after which *P*(*t*) > *P*\*, *Q*(*t*) > *Q*\*, *C*(*t*) := *P*(*t*) + *Q*(*t*) > *P*\* + *Q*\* =: *C*\* and *V*(*t*) > *V*\*, respectively. Simulation of the perturbed model highlighted that increasing *μ*_*c*_ causes all time thresholds 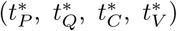 to increase, whereas increasing *μ*_*v*_ causes an increase in 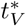 only (Supplementary Figure 28). Supplementary Figures 29 - 31 show how the time thresholds vary when all other model parameters vary and highlight that all of these thresholds are extremely sensitive to the values of *λ*, *μ*_*c*_, *γ*_*c*_ and *P*_0_.

## Discussion

By combining cell tracking methods with computational models we derived quantitative estimates of the proliferative activity of crypt stem cells and of their influence on villus cell migration during inflammatory conditions. Experimental time courses were analysed by fitting the data to compartmental models with two and three compartments. Both fitted models were able to reproduce well the trend of the experimental time courses. The three-compartment model allowed prediction of the time evolution of proliferative and non-proliferative cells at the expense of requiring estimation of a greater number of unknown parameter values when compared to the two-compartment model. The posterior parameter and predictive distributions highlighted in both models that, whereas an acute and temporary increase of inflammatory cell death did not influence distinctly net cell proliferation (new born cells minus dead cells) and migration onto the villus, a prolonged and less severe inflammation caused a decrease in net cell proliferation which produced, in turn, a delayed migration. To further investigate how injury may affect the dynamics of cells in the epithelium and trigger such delay, we simulated cell death, caused by inflammation, by means of a cell-based model and generated simulated time courses. Analysis of these time courses by means of compartmental models showed delayed migration under simulated chronic injury as experimentally observed, highlighting how a prolonged increase in cell death affects the dynamics of cells in the epithelium by delaying their migration. In summary, integration of computational modelling with experimental data derived from cell tracking methods allowed us to distinguish which inflammatory conditions influence epithelial cell dynamics. Identification of such conditions may highlight their contribution to barrier dysfunction in the development of intestinal inflammation. To the best of our knowledge, an experimental and computational analysis of cell dynamics during villus injury such as the one described in this article, which integrates compartmental and cell-based models with novel experimental time courses, has not been presented before. We believe that this analysis may stimulate further experimental work to estimate, for example, the proportion of proliferative and non-proliferative cells in the crypts.

## Data accessibility

The datasets supporting this article have been uploaded as part of the electronic supplementary material.

## Authors’ contribution

D. Muraro performed the computational analysis of the mathematical models; A. Parker and L. Vaux designed and performed the experiments; S. Filippi contributed to implementing the inference of the model parameters; A. G. Fletcher, P. K. Maini and H. M. Byrne contributed to designing the work and developing the mathematical models; A. J. M. Watson participated in designing the work and in experimental planning; C. Pin contributed to project design, mathematical model development, experimental planning and data analysis. All of the authors contributed to writing the manuscript.

